# RvD1 and LXA4 inhibitory effects on cardiac voltage-gated potassium channels

**DOI:** 10.1101/2024.10.29.620806

**Authors:** Alicia De la Cruz, Carlotta Ronchi, Chiara Bartolucci, Paula G Socuéllamos, Angela de Benito-Bueno, Stefano Severi, Antonio Zaza, Carmen Valenzuela

**Author notes:** Corresponding authors: Carmen Valenzuela. Instituto de Investigaciones Biomédicas Sols-Morreale CSIC-UAM. C/Arturo Duperier 4, 28029 Madrid, Spain. /4499. Alicia de la Cruz. Department of Biomedical and Clinical Sciences, Linköping University, Linköping, Sweden.

## Abstract

**Aims:** The resolution of inflammation is modulated by specialized pro-resolving lipid mediators (SPMs), which can be modified in some cardiovascular diseases. Among them, RvD1 and LXA_4_ prevent atrial fibrillation (AF) remodeling in the atria and cardiac hypertrophy, respectively in animal models. However, little is known about their electrophysiological effects on cardiac voltage-gated (VG) ion channels.

**Methods and results:** We used the patch-clamp technique in heterologous systems and cardiomyocytes to assess the acute effect of RvD1, and LXA_4_, on VG potassium currents. *In silico* simulations were used to predict the effect of current modulation on the atrial and ventricular action potentials (AP). RvD1 and LXA_4_ reduced *I*_Ks_ (channel K_V_7.1/KCNE1) in COS-7 cells and guinea-pig cardiomyocytes without modifying its voltage dependence; RvD1 was more potent than LXA_4_. In heterologous systems, RvD1 was also tested on *I*_Kur_ (channel K_V_1.5), *I*_to_ (channel K_V_4.3/KChIP2), *I*_Kr_ (channel K_V_11.1), and *I*_K1_ (channel K_ir_2.1) with the largest inhibitory effect on *I*_Ks_ and *I*_Kr_. In simulations RvD1 prolonged repolarization significantly in both atrial and ventricular myocytes.

**Conclusion:** The results provide a comprehensive evaluation of RvD1 and LXA_4_ on cardiac human potassium channels, at pathophysiological relevant concentrations, being RvD1 more potent than LXA_4_. The predicted effects on the action potential suggest that, along with their antiinflammatory action, RvD1 may reverse AF-induced electrical remodeling in the atria by direct modulation of K^+^ currents. The same action might instead contribute to ventricular functional remodeling; however, direct evidence for this is missing.

## 1. Introduction

Derivatives of n-3 and n-6 polyunsaturated fatty acids (PUFAs), collectively named SPMs, include several members grouped into 4 families: lipoxins, protectins, maresins, and resolvins.^1–4^ Among SPMs, resolvins (RvDs) may be endowed with antiarrhythmic effects; indeed, RvD1 prevented AF and countered atrial pro-arrhythmic remodeling in experimental models.^5,6^

Inflammation strongly contributes to the progression of cardiovascular disease^6^ and RvD1 is endowed with a potent anti-inflammatory and pro-resolution effectiveness.^7–10^ Moreover, RvD2 protected cardiovascular function and structure, by modulating vascular factors, fibrosis, and inflammation, when administered before and after the development of hypertension.^11^ Therefore, SPMs antiarrhythmic effects have been so far attributed to their role in the resolution of inflammation. Nonetheless, the effects of polyunsaturated fatty acids (PUFAs) (SPMs precursors) directly modulate voltage-gated (VG) ion channels (Na_V_, Ca_V_ and K_V_), to the extent that they have been proposed as potential anti-arrhythmic drugs.^8,12–19^ Therefore, direct regulation of ion channels by SPMs, thus far poorly investigated, might be relevant to cardiac electrophysiology.

The present study describes the effect of resolvins (RvD1), and lipoxins (LXA_4_) on a variety of VG K^+^ channels, an information relevant to the pathogenesis of arrhythmias and evolution of cardiac diseases, whether including inflammation or not. Our results show that RvD1 and LXA_4_ inhibited K_V_7.1/KCNE1 and K_V_11.1 channels, with some changes in their voltage dependence, being RvD1 more effective than LXA_4_. K_V_1.5, K_ir_2.1, and K_V_4.3/KChIP2 channels were affected by SPMs to a lesser extent.

## 2. Materials and Methods

### 2.1. Experimental models

#### 2.1.1. Primary cells. Guinea-pig cardiac ventricular myocytes

All animal care and experimental procedures were performed conforming to the NIH guidelines (Guide for the care and use of laboratory animals) revised in 2011; and from Directive 2010/63/EU of the European Parliament on the protection of animals used for scientific purposes and approved by the University of Milano-Bicocca ethics review board.

Adult Dunkin-Hartley Guinea-Pig (body weight 300 g, Envigo) were anesthetized with Ketamine + Xylazine (130 +7.5 mg/Kg) i.p. and euthanized by cervical dislocation.

#### 2.1.2. Isolation of cardiomyocytes, culture of cell lines and transfection

Ventricular cardiomyocytes were isolated by using a previously published^20^ retrograde aortic perfusion method with minor modifications. After isolation, the medium Ca^2+^ concentration was gradually restored to 1 mM and the cells were stored at +4°C until use. Only rod-shaped, Ca^2+^- tolerant cardiomyocytes were used within 12h from dissociation. Electrophysiological measurements in guinea-pig ventricular cardiomyocytes were performed with the whole-cell patch-clamp configuration.

COS-7, *Ltk^-^* and CHO-K1 cells were cultured in DMEM (COS-7 and *Ltk^-^*) or Iscove’s modified Eagle’s (CHO-K1) supplemented with 10% (v/v) FBS, and 100 IU/ml of penicillin and 100 µg/ml streptomycin (all from Gibco).^14^ CHO-K1 cells medium was supplemented with 1% (v/v) L-Glutamine; and *Ltk^-^* cells medium was supplemented with 0.25 mg/ml G418. Cells were cultured in an atmosphere with 5% CO_2_ at 37°C. COS-7 cells have an endogenous acid-sensitive potassium current,^21^ which did not interfere with the recordings obtained after transfecting the cells with K_V_7 or K_V_1 subfamilies channels nor with K_ir_2.1, which were the potassium channels of interest to us.

For electrophysiological recordings, COS-7 cells were incubated in 35 mm pretreated sterilized culture dishes (Falcon) and transiently transfected with 0.5 µg pEYFP-N1-KCNQ1 or co-transfected with 0.5 µg pEYFP-N1-KCNQ1 and 0.5 µg pECFP-C1-KCNE1. To record *I*_K1_, 0.5 µg of K_ir_2.1 cloned into pcDNA3.1 (gift from Dr. E. Delpón, Universidad Complutense de Madrid, Spain) was transfected in COS-7 cells for electrophysiological recordings. Electrophysiological recordings of K_V_11.1 were carried out in COS-7 cells after transfection with 1 µg of K_V_11.1 (cloned into a pcDNA3.1, kindly provided by Drs. S. Nattel, T. E. Hébert, and W. Weerapura). Stably transfected *Ltk^−^* cell line with the gene encoding K_V_1.5 channel was a gift of Dr. M.M. Tamkun (Colorado State University, USA).^22^ Before use, subconfluent cultures were incubated with 2 μM dexamethasone (D4902, Sigma-Aldrich, Saint Louis, MO, USA) for 24 h. CHO-K1 cells were transiently co-transfected with 2 µg K_V_4.3 (cloned into pBK-CMV vector, amplified by PCR and ligated into pmCherry-C1 (Clontech), by using *XhoI* and *HindIII* restriction sites,^23^ and 1 µg KChIP2 (cloned into IRES mCherry KChIP2 using *XbaI* and *XhoI* restriction sites).

In all cases, the cDNA were transient transfected together with 0.5 µg of the reporter plasmid EBO-pcD-Leu2-CD8 (to select transfected cells) per 35 mm culture dish^24^ in cells at 60-80% confluence following the FuGENE6 transfection method (Promega). The ratio µg DNA:µl Fugene was 1:3. Before use, cells (COS-7 and CHO-K1 cells) were incubated with polystyrene microbeads precoated with anti-CD8 antibody (Dynabeads M450, Dynal) and were removed from culture plates using TrypLE™ Express (Life Technologies).^14, 25^

### 2.2. Electrophysiological recordings

Guinea-pig ventricular cardiomyocytes were superfused through a thermostated pipette; the solution temperature was also monitored at the pipette tip with a fast-response digital thermometer (BAT-12, Physitemp, Clifton, NJ, USA) and kept at 36 ± 0.5°C. The extracellular (bath) solution contained (in mM): 154 NaCl, 4 KCl, 2 CaCl_2_, 1.2 MgCl_2_, 5 HEPES-NaOH, and 5.5 glucose (pH=7.4 with KOH); and intracellular (pipette-filing) solution contained (in mM): 110 K-aspartate, 23 KCl, 2 CaCl_2,_ 5 phosphocreatine, 3 MgCl_2_, 0.4 Na-GTP, 5 Na-ATP, 5 HEPES-K, and 5 EGTA-K (pH=7.3 with KOH). Membrane capacitance and series resistance were measured in every cell. Signals were filtered at 0.4 kHz (Axon Multiclamp 700B, Molecular Devices, Sunnydale, CA, USA), and digitized through a 12-bit A/D converter (Axon Digidata 1440A, Molecular Devices, Sunnydale, CA, USA) with a sampling rate of 1 kHz. The slow delayed rectifier potassium current (*I*_Ks_) was isolated as the current sensitive to 2 µM JNJ303 (Tocris, Bristol, UK). 10 µM nifedipine 5 µM E4031 and 1 µM SEA0400 (all from Sigma-Aldrich, Saint Louis, MO, USA), were added to Tyrode’s solution to minimize contamination by the L-type Ca channel (*I*_CaL_), the rapid delayed rectifier potassium current (*I*_Kr_), and the Na^+^/Ca^2+^ exchanger current respectively. The Na^+^ current was inactivated by setting the holding potential at −40 mV. *I*_Ks_ was activated by 5 sec V-steps from a holding potential of −40 mV. In Protocol 1 *I*_Ks_ was measured at a single potential (+40 mV) before and after steady-state drug application within the same myocyte (internal control); in Protocol 2, complete *I*_Ks_ current-voltage (I-V) relationships were obtained in separate cell groups previously incubated with control and SPMs-containing solutions respectively. *I*_Ks_ amplitude was measured at the end of the activating step (end-step current) and immediately after the return to the holding potential (tail current). To obtain the activation curves, tail current amplitudes were normalized to the maximal one.

For measurements in transfected cell lines (COS-7, CHO-K1 and *Ltk^-^*cells), the external solution contained (in mM): 130 NaCl, 4 KCl, 1.8 CaCl_2_, 1 MgCl_2_, 10 HEPES-Na, and 10 glucose (pH=7.4 with NaOH). Internal pipette-filing solution contained (in mM): 80 K-aspartate, 50 KCl, 3 phosphocreatine, 10 KH_2_PO_4_, 3 Mg-ATP, 10 HEPES-K, and 5 EGTA-K (pH=7.25 with KOH). *I*_Kur_ (channel K_V_1.5), *I*_to_ (channel K_V_4.3/KChIP2), *I*_Kr_ (channel K_V_11.1), and *I*_K1_ (channel K_ir_2.1) were recorded in the whole-cell configuration as previously reported^26^ at room temperature (23-25°C). *I*_Ks_ (from K_V_7.1 channels alone, or coexpressed with KCNE1) was recorded in the perforated patch-clamp configuration (Amphotericin-B, disolved in DMSO, was added to the pipette solution to achieve a final concentration of 100 μg/ml).^14^ I-V relationships were constructed by measuring the currents at the end of the activating step.^27^

Micropipettes were pulled from borosilicate glass capillary tubes (GD-1 model, Narishige, London, UK) on a horizontal puller (P-87, Sutter Instrument Co., CA, USA) and heat-polished with a microforge (MF-83, Narishige). After heat-polishing, the resistance of the patch electrodes tip (filled with the internal solution) averaged 2-4 MΩ.^14^ GΩ seal formation was achieved by suction; thereafter, capacitance and series resistance were compensated by about 80%. As a result, uncompensated series resistances were in the 4-8 MΩ range, thus, biasing voltage by less than 2 mV.

Signals were amplified by Axopatch-200B (Molecular Devices, Sunnydale, CA, USA), lowpass filtered (1 kHz) and sampled at 2 kHz (*I*_Ks_) or 4 kHz (other currents) by the A/D converter Digidata 1440A (Molecular Devices). Signals were further filtered (4-pole Bessel filter) at 1 kHz (*I*_Ks_) or 2 kHz (other currents). Command voltages and data acquisition were controlled with pClamp11 software (Molecular Devices).

The holding potential was set to −80 mV (unless otherwise indicated), and the interpulse interval was set to 10 s or 30 s, depending on the ion current recorded. The voltage pulse protocols were designed to analyze the biophysical properties of each channel. Time constants of current activation, inactivation, and deactivation were determined by fitting the recordings with mono- or bi-exponential functions:

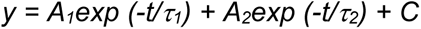

where τ_1_ and τ_2_ are time constants, A_1_ and A_2_ are the amplitudes of each component of the exponential, and C is the baseline value.^14,24,26,27^ The voltage dependence of channel activation was fitted to a Boltzmann equation:

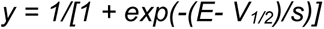

in which *s* represents the slope factor, E represents the applied voltage, and *V_1/2_* the voltage at which 50% of the channels are activated.^24,28^ Origin 2020b (OriginLab Software, Northampton, MA, USA) and CLAMPFIT 10.6 software from the PCLAMP 10 package were used to analyze data, perform least-squares fitting and for data presentation.

### 2.3. In-silico action potential analysis

To predict the effects of RvD1 on the action potential (AP), two human ventricular models, the Bartolucci-Passini-Severi (BPS)^29^ and the O’Hara–Rudy (ORd),^30^ and two human atrial models Mazhar-Bartolucci-Severi (MBS) and the Koivumaki (K)^31,32^ were used. For ventricular simulations, models for for isolated endocardial, epicardial, and M myocytes were used. To simulate RvD1 (at 5 and 50 nM) effects, the models were parametrized according to the experimental results reported in Table 1. The effect on activation V-dependency was simulated by changing the original V_1/2_ parameter by the compound-induced shift (mV) and the slope factor by the % compound-induced change. Simulations were carried out by considering RvD1 effect on a single current at a time or by incorporating all the affected currents. All models were paced at the basic cycle length of 1000 ms to steady state (> 1000 cycles). The action potential duration was measured at 90% repolarization (APD_90_). Model equations were implemented in Matlab 2021b (The MathWorks Inc., Natick, MA, USA) and numerically solved using the ode15s solver.

**Table 1.** Effects of SPMs on the voltage dependence on K_V_7.1/KCNE1, K_V_7.1, K_V_1.5 and K_V_4.3/KChIP2 and K_V_11.1.

|  | K<sub>V</sub>7.1/KCNE1 |  | K<sub>V</sub>7.1 |  | K<sub>V</sub>1.5 |  | K<sub>V</sub>4.3/KChIP2 |  | K<sub>V</sub>11.1 |  |
| --- | --- | --- | --- | --- | --- | --- | --- | --- | --- | --- |
|  | V<sub>1/2</sub> (mV) | s (mV) | V<sub>1/2</sub> (mV) | s (mV) | V<sub>1/2</sub> (mV) | s (mV) | V<sub>1/2</sub> (mV) | s (mV) | V<sub>1/2</sub> (mV) | s (mV) |
| **Control** | 34.5±4.9 | 20.2±1.8 | -16.5±2.3 | 7.1±0.8 | -12.9±1.0 | 5.7±0.4 | 26.3±2.4 | 15.4±1.1 | -6.6±2.3 | 8.3±0.4 |
| **RvD1**<br>**(5 nM)** | 35.6±11.1 | 17.9±1.6 | -11.6±1.8* | 8.9±0.7 | -14.3±1.3 | 6.1±0.4 | 34.0±1.9* | 17.4±0.5 | -7.5±4.5 | 8.5±0.4 |
| **Control** |  |  |  |  | -11.5±1.4 | 5.6±0.4 | 26.3±2.4 | 15.4±1.1 | -6.5±3.8 | 8.6±0.5 |
| **RvD1**<br>**(50 nM)** |  |  |  |  | -10.0±2.3 | 5.1±0.6 | 29.7±2.3* | 16.4±1.0 | -2.7±4.5* | 7.0±0.9 |
| **Control** | 48.5±2.4 | 21.8±2.2 |  |  |  |  |  |  |  |  |
| **LXA<sub>4</sub>**<br>**(500 nM)** | 42.0±4.1 | 21.7±1.3 |  |  |  |  |  |  |  |  |

Data are shown as the mean ± SEM. \*: $P < 0.05$ (Paired Student's t-test), n = 4-8.

### 2.4. Drugs and reagents

(S)-Resolvin D1 (or RvD1) (Cayman Chemical, Ann Arbor, MI, USA), and LXA_4_ (Santa Cruz Biotechnology, Dallas, USA) were dissolved in ethanol (final concentration 1 μM) and stored protected from light at −80°C. For electrophysiological recordings, each SPM was dissolved in the extracellular solution at concentrations in the 5-500 nM range. 0.5-500% ethanol was added to the control solutions.

### 2.5. Statistical analysis

Data are expressed as mean ± SEM of n measurements. Comparisons between mean values were performed by using the Student’s t-test or ANOVA, as appropriate (Bonferoni’s correction in post–hoc comparisons). Statistical significance was defined as *P* < 0.05. Sample size (n) is reported in figure legends.

## 3. Results

### 3.1. RvD1 and LXA_4_ effects on K_V_7.1/KCNE1 in COS-7 cells

Figure 1A shows *I*_Ks_ (K_V_7.1/KCNE1) recordings, in the absence and the presence of RvD1. RvD1 5 and 50 nM inhibited *I*_Ks_ amplitude by 48.6 ± 5.7 % (n = 5, *P* < 0.01) and 86.3 ± 5.0 % (n = 6, *P* < 0.01) respectively, measured at the end of 5.5 s pulses at +60 mV.

**Figure 1:**
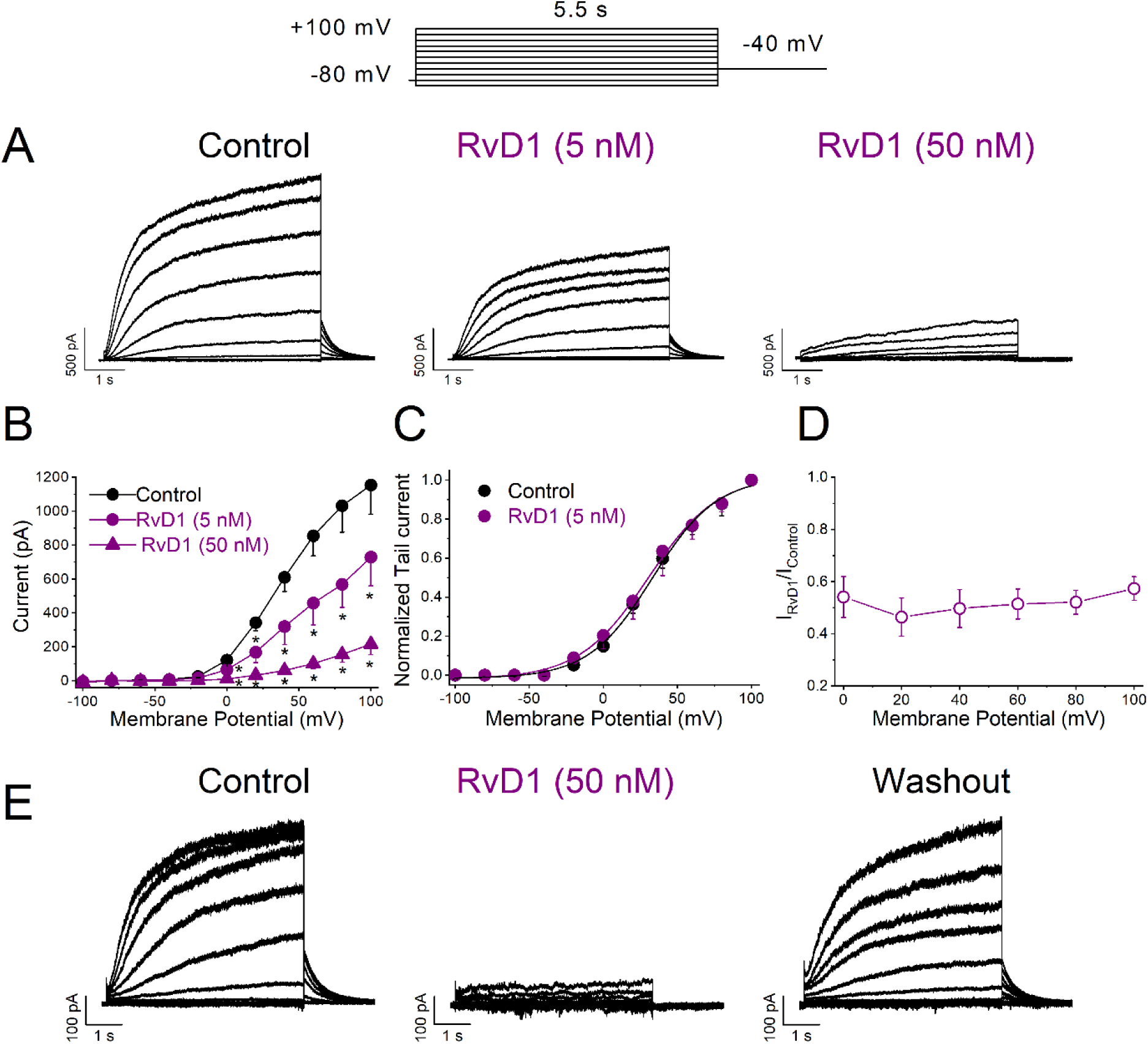
Voltage-dependent effects of RvD1 on K_V_7.1/KCNE1 (*I*_Ks_) in transiently transfected COS-7 cells. A: Current traces in the absence and in the presence of 5 and 50 nM RvD1. B: I-V relationships obtained after plotting the current at the end of 5.5-s vs. membrane potential, in the absence and in the presence of RvD1. C: Activation curves of *I*_Ks_ in the absence and in the presence of 5 nM RvD1. D: Ratio between the current in the presence of 5 nM RvD1 and under control conditions. E: Representative traces of 50 nM RvD1 wash-out on *I*_Ks_. Data are shown as the mean ± SEM. \**P* < 0.05 (Paired Student’s t-test), n = 5-8.

Figure 1B shows the RvD1 (5 and 50 nM) effect on *I*_Ks_ I-V relationship. At both concentrations, RvD1 inhibited *I*_Ks_ at membrane potentials positive to 0 mV. Neither the activation V_1/2_ nor the slope factor were modified (Table 1). RvD1 effects were neither voltage-dependent, (Figure 1D), nor time-dependent (Tables 2 and 3). *I*_Ks_ inhibition by RvD1 was largely reversible upon wash-out (73.5 ± 17.8 %, n = 4 vs. control conditions, *P* > 0.05, Figure 1E).

**Table 2.** Activation Kinetics.

| **Kv7.1/KCNE1** |  |  |  |  |
| --- | --- | --- | --- | --- |
| *Control* |  | *RvD1 (5 nM)* |  |  |
| $\tau_s$ (ms) | $\tau_f$ (ms) | $\tau_s$ (ms) | $\tau_f$ (ms) | (n, P) |
| 3608.3 ± 981.4 | 908.5 ± 197.1 | 3577.5 ± 784.1 | 920.2 ± 181.4 | n = 6,<br>P > 0.05 |
| *Control* |  | *LXA<sub>4</sub> (500 nM)* |  |  |
| $\tau_s$ (ms) | $\tau_f$ (ms) | $\tau_s$ (ms) | $\tau_f$ (ms) | |
| 3340.0 ± 851.7 | 726.5 ± 160.4 | 2506.7 ± 369.8 | 674.5 ± 194.4 | n = 7,<br>P > 0.05 |
| **Kv7.1** |  |  |  |  |
| *Control*<br>$\tau$ (ms) | | *RvD1 (5 nM)*<br>$\tau$ (ms) | | |
| 16.0 ± 4.5 | | 34.7 ± 9.2 | | n = 3,<br>P > 0.05 |

Data are shown as the mean ± SEM. \*: P < 0.05 (Paired Student's t-test), n = 3-7.

RvD1 (5 and 50 nM) effects on K_V_7.1 in the absence of KCNE1 were also studied (Figure S1). At 5 nM, RvD1-induced block was weaker than that observed with this compound on K_V_7.1/KCNE1 (28.9 ± 5.8 % n = 5, *P* < 0.05, measured at the end of 5.5 s pulses at +60 mV). However, in this case, RvD1 shifted the activation V_1/2_, by + 5mV without modifying the slope factor (Table 1) and activation kinetics (Tables 2 and 3). These results suggest that KCNE1 tunes the sensitivity of K_V_7.1 to RvD1, being RvD1 more potent to block these channels in the presence of this β-subunit (Figures 1 and S1).

Figure 2 shows the effects of 500 nM LXA_4_ on *I*_Ks_. LXA_4_ inhibited *I*_Ks_ by 33.0 ± 3.4 % (n= 5, *P* < 0.01) measured at the end of 5.5 s pulses at +60 mV, (Figure 2A). As shown by the I-V relationship (Figure 2B), LXA_4_ decreased the current magnitude at membrane potentials positive to 0 mV.

**Figure 2:**
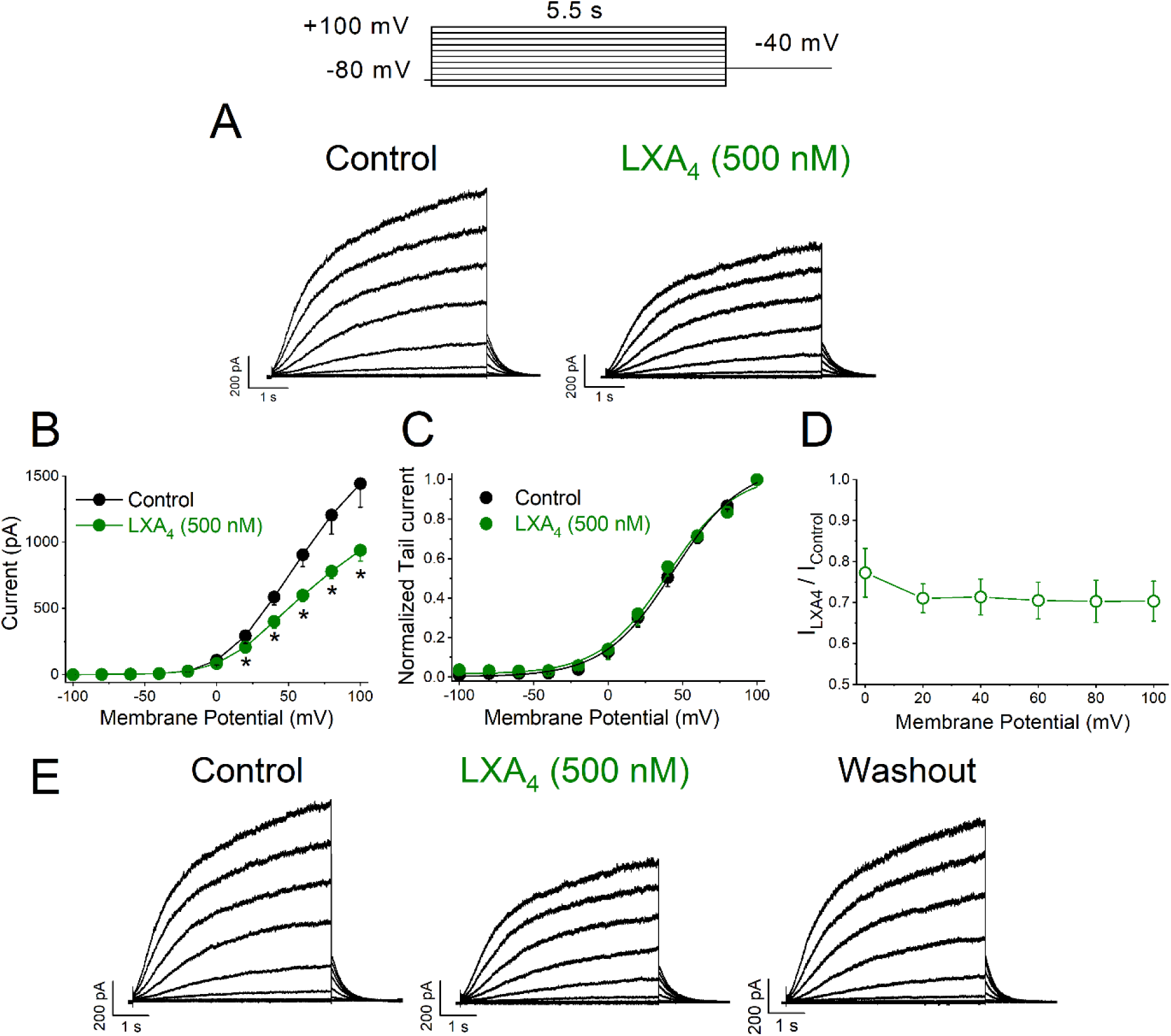
Voltage-dependent effects of LXA_4_ on K_V_7.1/KCNE1 (*I*_Ks_) in transiently transfected COS-7 cells. A: Current traces were obtained after applying the pulse protocol shown in the top, both in the absence and in the presence of LXA_4_ (500 nM). B: I-V relationships obtained under control conditions and after perfusion with LXA_4_. C: Activation curves of *I*_Ks_ obtained after plotting the maximum tail current amplitude vs. the previous step potential in the absence and in the presence of LXA_4_. D: Ratio between the current in the presence and in the absence of LXA_4_. E: Representative traces of LXA_4_ wash-out on *I*_Ks_. Data are shown as the mean ± SEM. *: *P* < 0.05 (Paired Student’s t-test), n= 5.

LXA_4_ modified neither the V_1/2_ nor the slope of the *I*_Ks_ activation curve (Figure 2C and Table 1). LXA_4_-induced block was not voltage-dependent, being similar at +20 mV and +100 mV (Figure 2D) and did not affect activation kinetics (Table 2). However, LXA_4_ slowed the deactivation process (Table 3). LXA_4_ effects were almost completely reversible on wash-out (89.3 ± 4.5%, n = 6, P > 0.05 vs. control conditions, Figure 2E). Overall, albeit qualitatively similar to that of RvD1, LXA_4_ effect on *I*_Ks_ was quantitatively smaller. Both agents failed to modify current kinetics and their effct was largely reversible.

**Table 3.** Deactivation Kinetics.

| **Kv7.1/KCNE1** |  |  |
| --- | --- | --- |
| *Control*<br>$\tau$ (ms) | *RvD1 (5 nM)*<br>$\tau$ (ms) | (n, P) |
| 426.1 ± 95.3 | 464.6 ± 90.6 | n = 5,<br>P > 0.05 |
| *Control*<br>$\tau$ (ms) | *LXA<sub>4</sub> (500 nM)*<br>$\tau$ (ms) | |
| 357.9 ± 50.1 | 429.6 ± 41.7* | n = 7,<br>P < 0.05 |
| **Kv7.1** |  |  |
| *Control*<br>$\tau$ (ms) | *RvD1 (5 nM)*<br>$\tau$ (ms) | |
| 303.0 ± 52.8 | 375.4 ± 92.3 | n = 3,<br>P > 0.05 |

Data are shown as the mean ± SEM. \*: P < 0.05 (Paired Student's t-test), n = 3-7.

### 3.2. RvD1 and LXA_4_ effects on *I*_Ks_ of guinea-pig ventricular cardiomyocytes

When using the internal control (Protocol 1) (Figure S2C), 5 nM RvD1 inhibited *I*_Ks_ measured at the end step by 29.9 ± 5.4 % at +40 mV (n ≥ 6, *P* < 0.05), the effect with 50 nM RvD1 was similar (28.1 ± 4.3%, n ≥ 6, *P* < 0.05). At the same potential, LXA_4_ (500 nM) inhibited the end-step of the *I*_Ks_ by 56.0 ± 6.0 % and the tail current by 45.2 ± 6.0% (n = 8, *P* < 0.05) (Figure S3B).

RvD1 and LXA_4_ effect on the *I*_Ks_ I-V relationship was tested by comparing separate cell groups incubated with the control and test solutions respectively (Protocol 2). Figure 3A shows representative recordings from three groups of cardiomyocytes in control conditions (black), after incubation with 5 nM RvD1 (red) and 50 nM RvD1 (blue) respectively. RvD1 5 nM significantly inhibited the end-step and tail currents (Figure 3B-C) at all potentials (n ≥ 10, *P* < 0.05); the step/tail ratio (at +60 mV) was unchanged (3.64 ± 0.57 vs 2.8 ± 0.16; NS). RvD 50 nM did not further reduce current amplitude, but significantly increased the step/tail raio (3.84 ± 0.35 vs 2.8 ± 0.16; *P* < 0.05), thus indicating an effect inversely proportional to the current driving force. Figure 3D shows that RvD1 (5 and 50 nM) did not modify the V_1/2_ or the slope factor of the activation curve (V_1/2_ : Control 21.7 ± 2.4 mV; RvD1 (5 nM) 17.4 ± 2.4 mV; RvD1 (50 nM) 25.6 ± 1.6 mV; Slope: Control 11.2 ± 0.95 mV; RvD1 (5 nM) 8.54 ± 1.1 mV; RvD1 (50 nM) 10.0 ± 1.8 mV).

**Figure 3:**
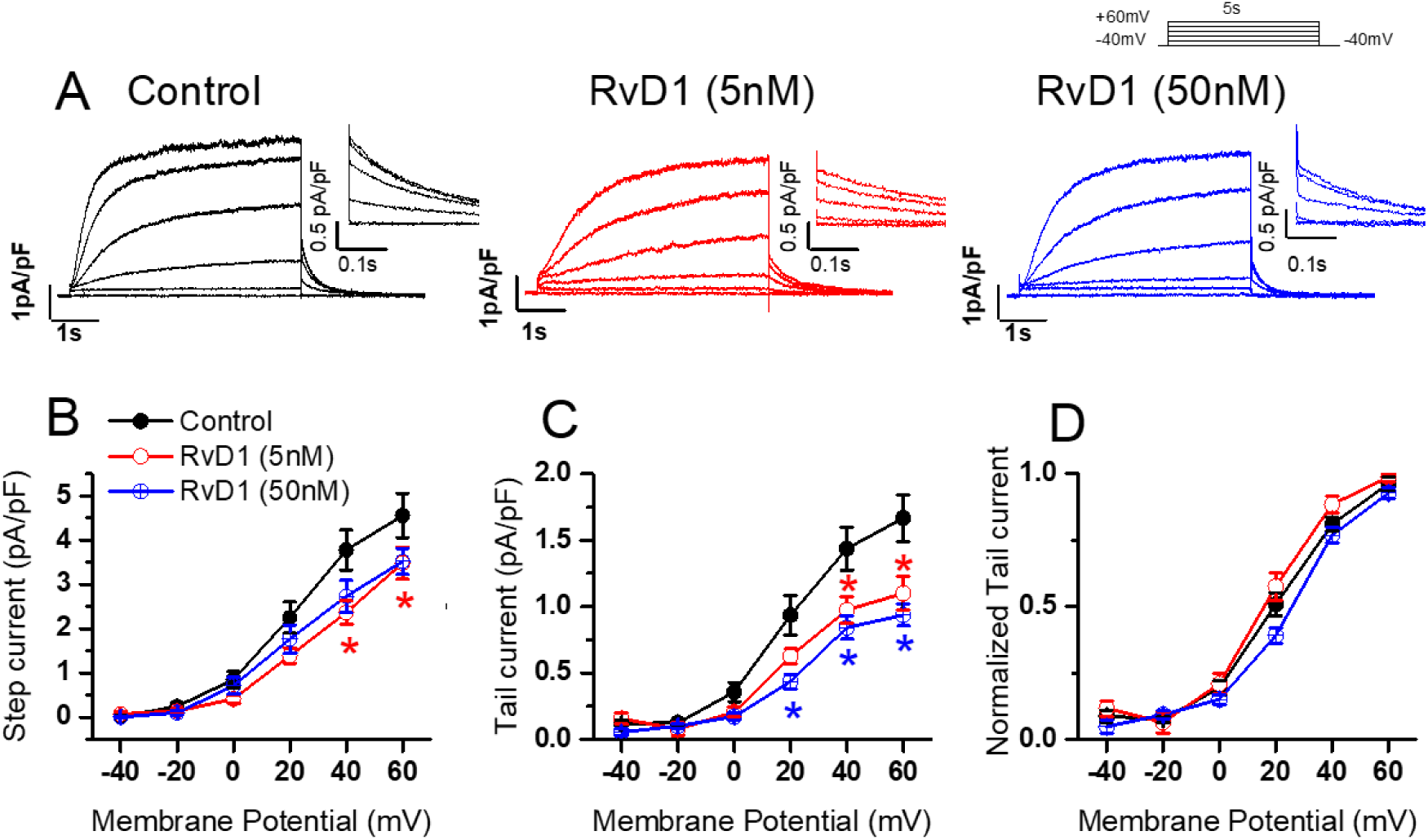
RvD1 inhibition on *I*_Ks_ of guinea-pig cardiomyocytes. A: Representative traces of *I*_Ks_ at various test potential in the 3 groups of myocytes: control (black), RvD1 5 nM (red) and RvD1 50 nM (blue); *I*_Ks_ tail currents magnified in the insets. B: Average (± SEM) I-V relationships for *I*_Ks_ at the end of the depolarizing step, C: *I*_Ks_ “tail current”. D: *I*_Ks_ voltage-dependent activation. n > 6 for all groups, * *P* < 0.05 vs. Control (Two way ANOVA).

Figure 4A shows representative *I*_Ks_ recordings from two groups of cardiomyocytes in control condition (black) and after incubation with control LXA_4_ (light blue). LXA_4_ inhibited the end-step and tail currents (Figure 4B-C) at all potentials, (n ≥ 10, *P* < 0.05); the step/tail ratio (at +60 mV) was unchanged (2.91 ± 0.57 vs 2.8 ± 0.16; NS). Figure 4D shows that LXA_4_ did not modify the *I*_Ks_ activation curve (V_1/2_ = 21.7 ± 2.4 mV vs. 15.7 ± 2.6, in control and LXA_4,_ respectively; Slope = 11.2 ± 0.95 mV vs. 9.5 ± 1.4 mV, in control and LXA_4,_ respectively; n = 15 and n = 8, in control and LXA_4,_ respectively, *P* > 0.05).

**Figure 4:**
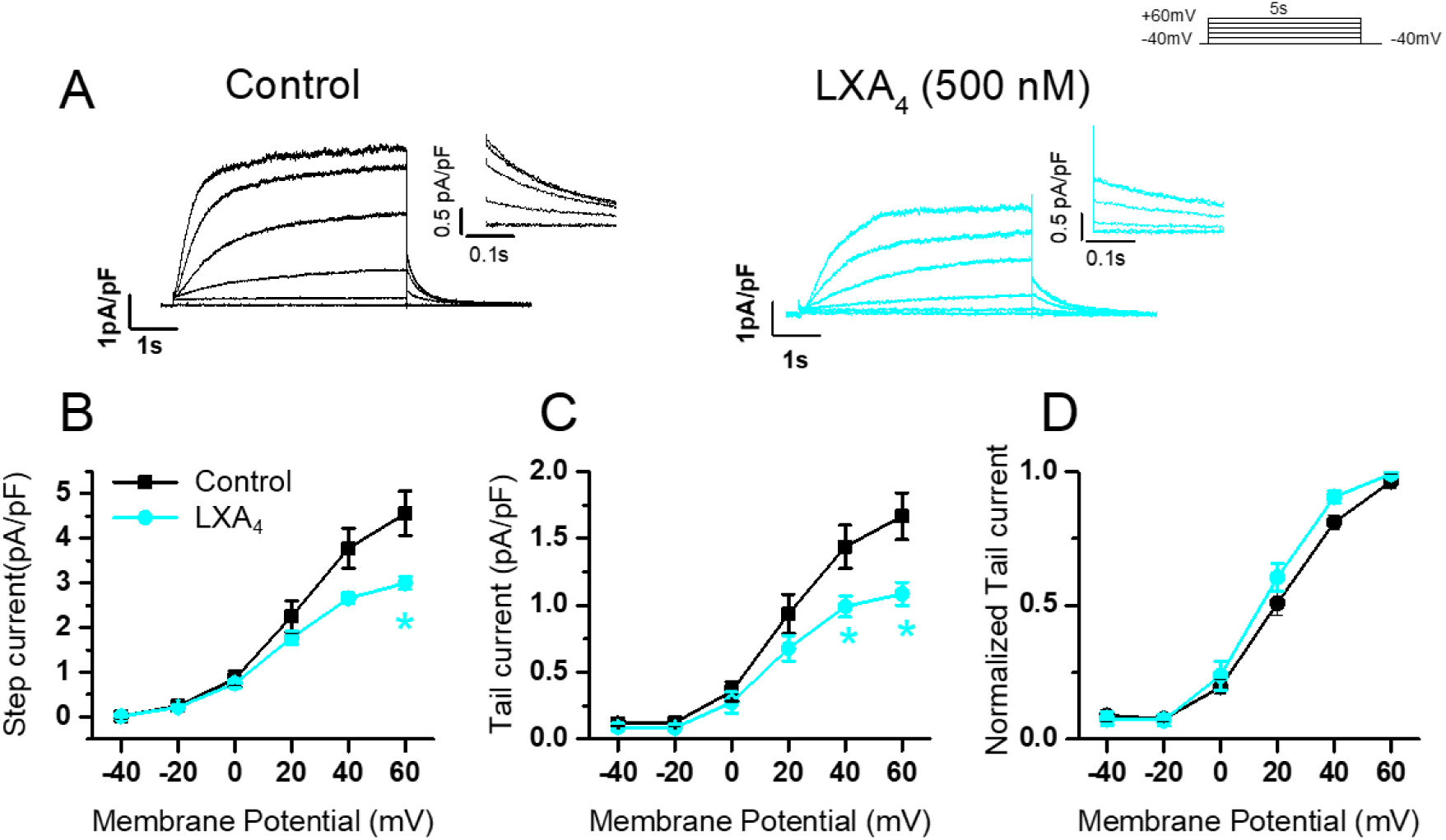
LXA4 inhibition on *I*_Ks_ of guinea-pig cardiomyocytes. Representative traces of *I*_Ks_ at various test potentials in the 2 groups of myocytes: control (black) and 500 nM (light blue). A: *I*_Ks_ tail currents magnified in the insets. B: Average (± SEM) I-V relationships for *I*_Ks_ at the end of the depolarizing step and C: *I*_Ks_ “tail current” in control and LXA_4_ group. D: *I*_Ks_ voltage-dependent activation in the two experimental groups. Control n > 10, LXA_4_ n > 6. * *P* < 0.05 vs. Control (Two way ANOVA).

### 3.3. RvD1 effects on other cardiac voltage-gated potassium channels

Next, we wanted to elucidate whether RvD1 also inhibits other cardiac potassium channels. To this aim, we studied the effects of RvD1 in heterologous systems (COS-7, CHO-K1 and *Ltk*^-^ cells) transfected with different cardiac potassium channels.

#### 3.3.1. RvD1 effects on K_V_1.5 (in *Ltk^-^* cells)

K_V_1.5 channels carry an ultra-rapid delayed rectifier potassium current (*I*_Kur_), which plays an essential role in phases 1 and 2 of atrial repolarization. Figure 5A-B show *I*_Kur_ and its I-V relationship. RvD1 (5 nM) slightly inhibited *I*_Kur_ (at the end of the activating step) at all potentials (at +60 mV 6.9 ± 1.0 %; n= 8, *P* < 0.01); and at 50 nM had a somewhat larger effect (28.2 ± 2.5 %; n = 7, *P* < 0.05) measured at the end of 250 ms at the same potential. The activation curve and the slope factor were not modified at either of the two concentrations tested (Table 1, Figure 5C). RvD1-induced block was not voltage-dependent (Figure 5D).

**Figure 5:**
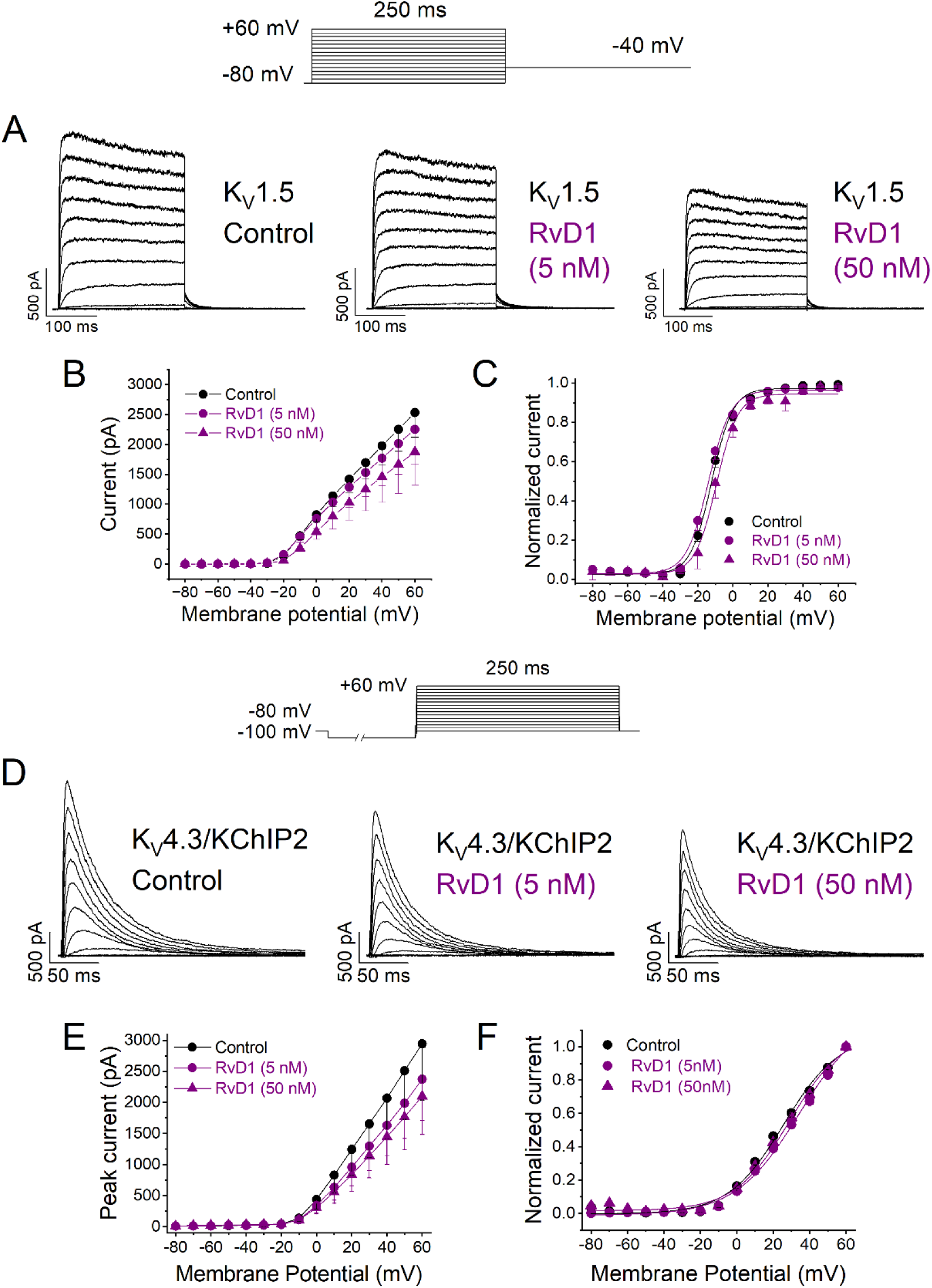
Effects of RvD1 on K_V_1.5 (*I*_Kur_) and K_V_4.3/KChIP2 (*I*_to_) in stable transfected *Ltk^-^* cells and transiently transfected CHO cells, respectively. Currents were elicited after applying the protocol shown on the top of the figure. Representative traces of *I*_Kur_ (A) and *I*_to_ (D) currents in *Ltk^-^* and CHO cells recorded in the absence (control) and in the presence of 5 and 50 nM RvD1. B: I-V *I*_Kur_ relationships obtained after plotting the current at the end of 250 ms vs. membrane potential, in the absence and the presence of RvD1. C: Activation curves of *I*_Kur_ in the absence and the presence of RvD1. E: I-V relationships were obtained after plotting the current measured at the maximum peak for every membrane potential tested vs. membrane potential, in the absence and the presence of RvD1. F: Activation curves of *I*_to_ in the absence and in the presence of RvD1. Data are shown as the mean ± SEM. *: *P* < 0.05 (Paired Student’s t-test), n = 4-8 per group.

#### 3.3.2. RvD1 effects on K_V_4.3/KChIP2 (in CHO cells)

The K_V_4.3/KChIP2 complex carries the main component of *I*_to_, the fast repolarizing current responsible for phase 1 of the AP, particularly prominent in atrial myocytes. Figures 5 E-F show that RvD1 5 nM moderately inhibited *I*_to_ (17.5 ± 2.2 % at +60 mV; n = 4, *P* > 0.05) at all potentials; RvD1 50 nM had a somewhat stronger effect (27.3 ± 2.0%; n = 4, *P* > 0.05). The activation V_1/2_ and the slope factor were not modified by 5 nM RvD1; and RvD1 50 nM slightly shifted V_1/2_ towards more positive potentials (Table 1, Figure 5G). Block produced by RvD1 was not voltage-dependent (Data not shown).

#### 3.3.3. RvD1 effects on K_V_11.1 (hERG) (COS-7 cells)

K_V_11.1 (hERG) channels carry *I*_Kr_, a major contributor to repolarization particularly in ventricular myocytes. *I*_Kr_ inactivates with membrane depolarization; thus, at positive step potentials, the current may be larger upon repolarization (*I*_tail_ current) than at steady-state during the activating step (*I*_SS_). The *I*_SS_/*I*_tail_ ratio quantifies *I*_Kr_ inactivation, a property pivotal in setting *I*_Kr_ profile during the AP.

RvD1 substantially inhibited *I*_Kr_ in a concentration-dependent fashion. Thus, for steps to −10 mV, *I*_ss_ was slightly inhibited by 21.8 ± 5.9 % (n = 4, *P* > 0.05) and 45.3 ± 9.4 % (n = 4, *P* > 0.05) at 5 and 50 nM RvD1, respectively (Figure S4). However, at the same step potential, *I*_tail_ was inhibited by 31.7 ± 4.6 % (n = 4, *P* < 0.05) and 56.1 ± 9.5 % (n = 4, *P* < 0.05) at 5 and 50 nM RvD1, respectively (Figures 6A-D and S4). Accordingly, RvD1 increased the *I*_ss_/*I*_tail_ significantly (Figure S4). RvD1 100 nM completely blocked *I*_Kr_ (Figure S6A). The *I*_Kr_ activation V_1/2_ and slope factor were not modified by 5 nM RvD1 (Table 1, Figure 6D). However, 50 nM RvD1 negatively shifted the midpoint of the activation curve (−6.2 ± 0.9 mV, P<0.05), while leaving the slope factor unchanged (Table 1, Figure 6D).

**Figure 6:**
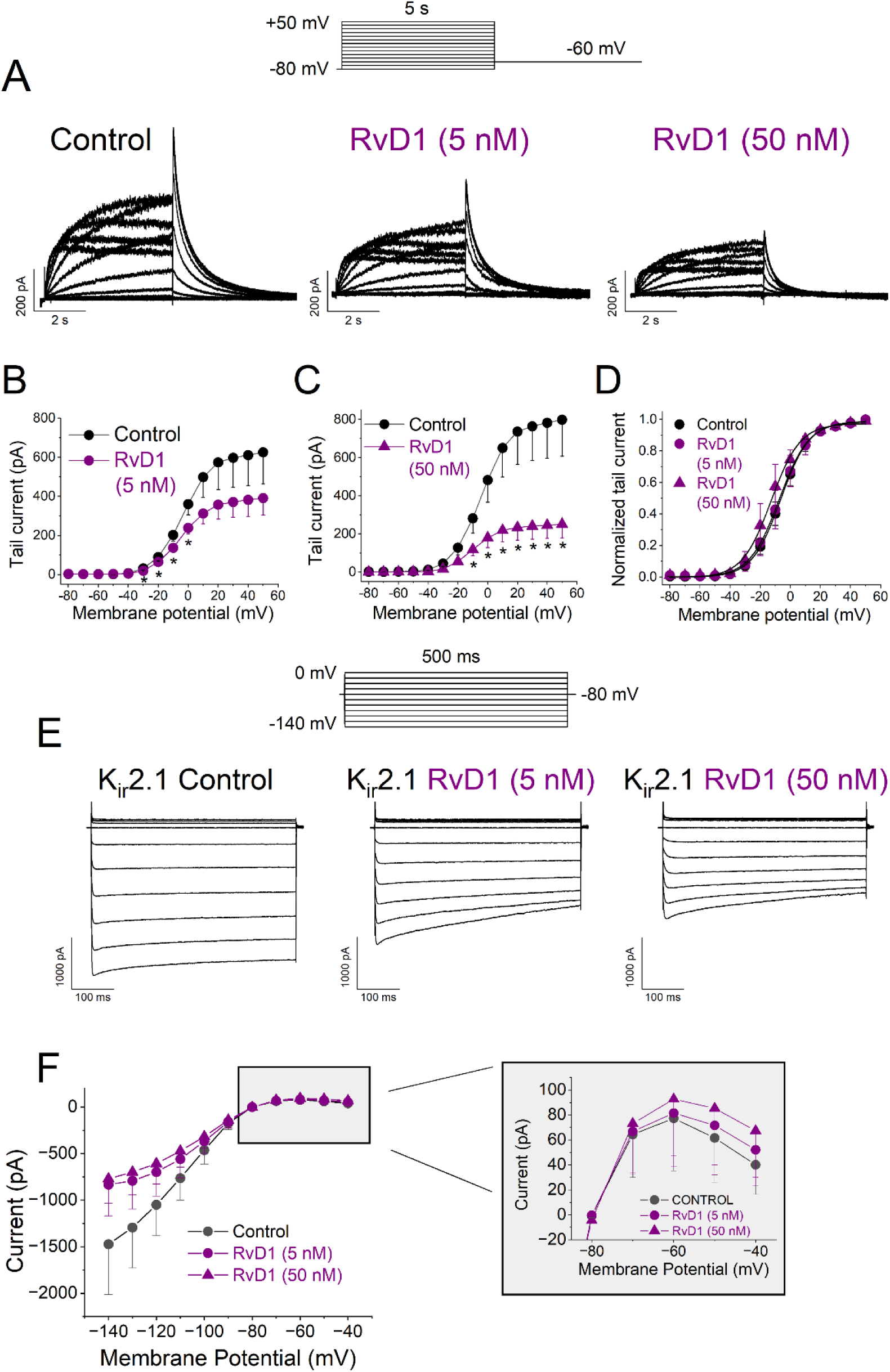
Effects of RvD1 on K_V_11.1 (*I*_Kr_) and K_ir_2.1 (*I*_K1_) in transiently transfected COS-7 cells. Currents were elicited after applying the protocol shown on the top of the figure (for *I*_Kr_ (A) and on top of original traces (for *I*_K1_, E). A: Representative traces of *I*_Kr_ recorded in the absence (control) and in the presence of 5 and 50 nM RvD1. B and C: *I*_Kr_ tail currents plotted vs. membrane potentials tested, in the absence and the presence of 5 nM RvD1 and 50 nM RvD1. D: Activation curves of *I*_Kr_ in the absence and the presence of RvD1. E: Representative traces of *I*_K1_ recorded in the absence (control) and in the presence of 5 and 50 nM RvD1. F: I- V relationships obtained after plotting the current at the end of 500 ms vs. membrane potential, in the absence and the presence of RvD1. The inset shows RvD1 effect on *I*_K1_ measured between −80 mV and −40 mV of the membrane potential. Data are shown as the mean ± SEM. *: *P* < 0.05 (Paired Student’s t-test), n = 4-5 per group.

#### 3.3.4. RvD1 effects on K_ir_2.1 (COS-7 cells)

K_ir_2.1 channels belong to the inward-rectifier potassium family and are expressed in excitable and non-excitable cells where they carry *I*_K1_.^33^ Outward *I*_K1_ is very small (strong inward rectification), thus steady-state inward *I*_K1_ was measured to quantify RvD1 effect. RvD1 5 and 50 nM similarly inhibited *I*_K1_. For instance, at −120 mV *I*_K1_ was reduced by 44.6 ± 2.0 % (n= 3, *P* < 0.05) and 46.9 ± 4.4 % (n = 4, *P* < 0.05) by 5 and 50 nM RvD1 respectively, measured at the end of 500 ms pulses. As shown by the I-V relationship, RvD1 blocked inward *I*_K1_ (at membrane potentialsbetween −140 and −70 mV), whereas outward *I*_K1_ was, if anything, slighly increased by RvD1 (Figure 6F). Inward *I*_K1_ tended to decrease over time during the voltage step, a pattern accentuated by RvD1 (Figure 6E); this might reflect a slow component of RvD1 interaction with the channel.

Figure S6 summarizes RvD1 effect on the channels analyzed in this study. It plots the percent current inhibition at the activation midpoint for each ion channel.

### 3.4. Computational modeling of RvD1 on cardiac AP

Numerical models were used to predict the effect of SPMs on ventricular (BPS model for endocardial, epicardial and M cells) and atrial (MBS model) AP morphology (Figure 7, Tables 4-5).

**Figure 7:**
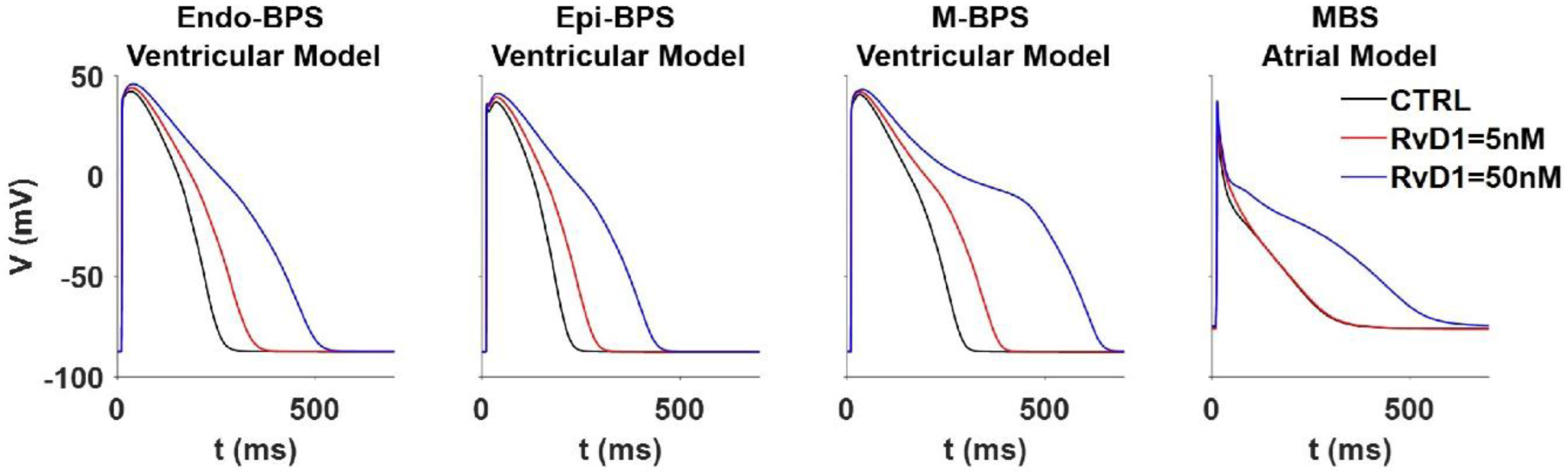
Computational modeling of RvD1 on cardiac AP. BPS ventricular cell computer model was used to examine the effects of RvD1, on AP morphology in A: endocardial, B: epicardial, and C: M cells. D: MBS human atrial AP model was used to examine the effects of RvD1, on atrial AP morphology.

**Table 4.**
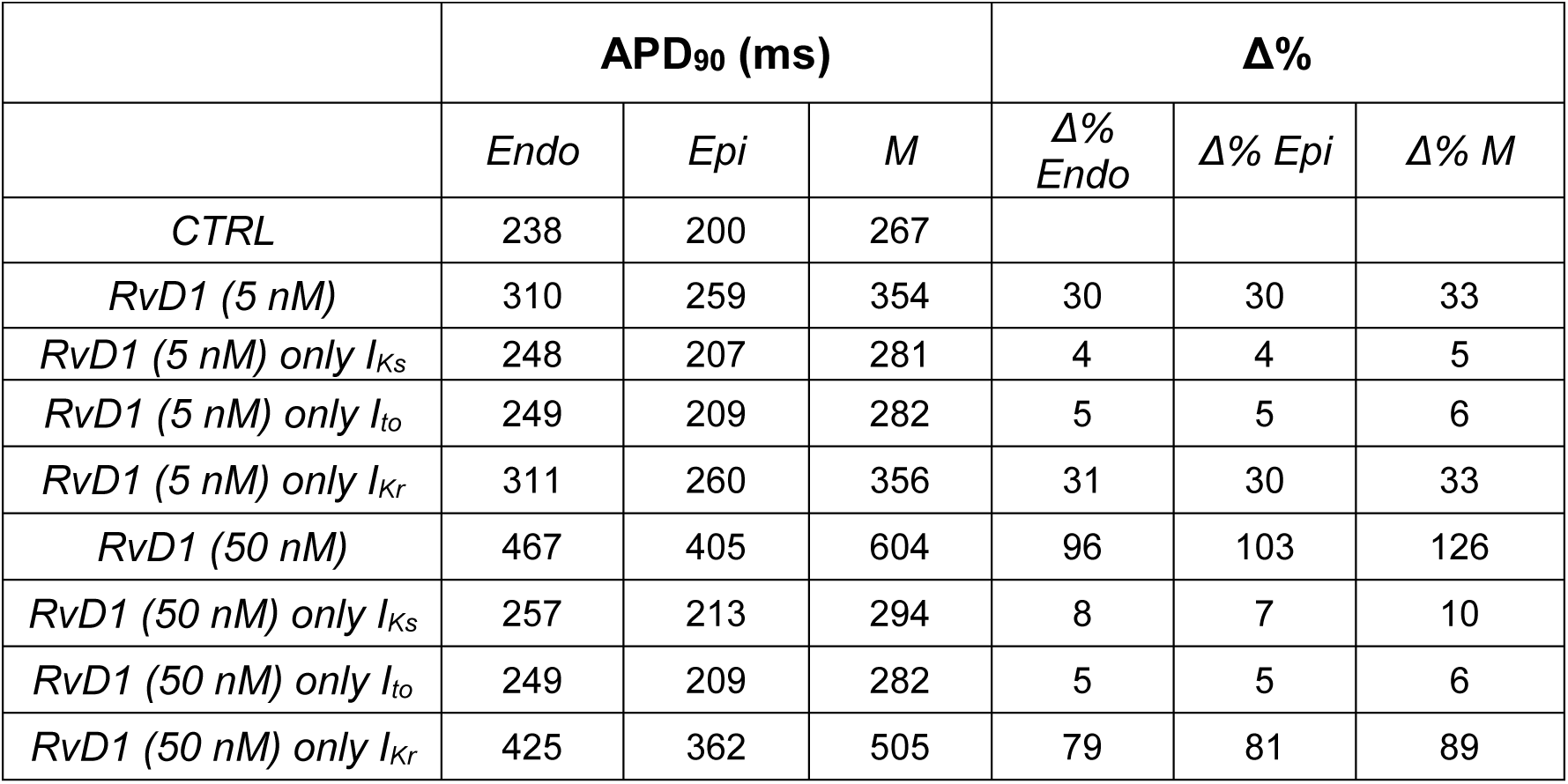
Computational modeling APD_90_ results with the BPS model.

**Table 5.** Computational modeling APD results with the atrial MBS model.

|  | **APD (ms)** | **Δ%** |
| --- | --- | --- |
| *CTRL* | 249 |  |
| *RvD1 (5 nM)* | 253 | 2 |
| *RvD1 (5 nM) only I<sub>Ks</sub>* | 258 | 3.6 |
| *RvD1 (5 nM) only I<sub>to</sub>* | 265 | 6.4 |
| *RvD1 (5 nM) only I<sub>Kr</sub>* | 259 | 4 |
| *RvD1 (5 nM) only I<sub>Kur</sub>* | 223 | -10 |
| *RvD1 (50 nM)* | 478 | 92 |
| *RvD1 (50 nM) only I<sub>Ks</sub>* | 264 | 6 |
| *RvD1 (50 nM) only I<sub>to</sub>* | 263 | 5.6 |
| *RvD1 (50 nM) only I<sub>Kr</sub>* | 270 | 8.4 |
| *RvD1 (50 nM) only I<sub>Kur</sub>* | 328 | 31.7 |

RvD1 dose-dependently prolonged APD in all types of ventricular myocytes (Figure 7). In atrial cells, APD prolongation was smaller and substantial only at 50 nM RvD1. In all cases, the main contribution to APD prolongation was given by *I*_Kr_ inhibition, as suggested by simulations in which only this current was modulated (see Table 4).

To rule out model dependency of the results, the simulations have been performed by using also the established ORd and K models and the AP behaviours are shown in Figure S5 and Tables S1-S2. Results are fully consistent with those obtained with the more recently developed BPS and MBS models. As only remarkable difference, in M cells, the effect of 50 nM of RvD1 was even exacerbated inducing membrane potential bistability and plateau-level oscillations (EADs) in M cells (Figure S5).

## 4. Discussion

This study analyzed the effects of two derivatives from n-3 PUFAs and n-6 PUFAs metabolism, RvD1 and LXA_4_ on VG cardiac potassium currents. The effect on *I*_Ks_ was consistent between ventricular myocytes and transfected cell-lines; only the latter were used to test the remaining currents. Our results demonstrate that both SPMs inhibit *I*_Ks_, but RvD1 effect was larger than that of LXA_4_. Noteworthy, RvD1 blocking effect on *I*_Ks_ was diminished when only the alpha-subunit of the K_V_7.1 channel was expressed, thus indicating that KCNE1 is also a target of RvD1. RvD1 also inhibited *I*_Kr_ (K_V_11.1 channel) substantially and dose-dependently, producing changes in the current rectifying behavior (the *I*_ss_/*I*_tail_ ratio) of potential relevance for the AP profile.

The potency of the RvD1 to inhibit K_V_1.5, K_V_4.3/KChIP2 and Kir2.1 channels was negligible in comparison to that observed on K_V_7.1/KCNE1 and K_V_11.1 channels. Finally, we used a holistic approach to predict RvD1 effects on cardiac action potentials by using different computational models. *In silico* results indicate that RvD1 prolongs the ventricular APD in a dose-dependent manner, mainly due to its inhibitory effect on *I*_Kr_ (Figure 7, Table 4).

### 4.1. Differences between experimental models and effect peculiarities

RvD1 and LXA_4_ inhibited *I*_Ks_ both in native guinea-pig myocytes and in a heterologous expression system (COS-7 cells); however, the magnitude of effects differed between the experimental models. Indeed, whereas in myocytes RvD1 effects on *I*_Ks_ was relatively small and saturated already at the lower concentration, a larger and dose-dependent effect was observed in COS-7 cells. Furthermore, at the same concentration (500 nM) LXA_4_ effect was larger in myocytes than in COS-7 cells. These discrepancies might reflect: i) a different response to SPMs of human (COS-7 cells) vs guinea-pig channel, ii) dependency of the response on “channelosome” proteins, available in the native myocytes but not in the heterologous system, or, iii) the effects produced in cardiomyocytes might be partially due to SPMs interaction with their membrane receptor (GPR32 or ALX/FPR2)^5^ not expressed in the cell line.

In COS-7 cells transfected with K_V_7.1 only, RvD1 modified the voltage dependence of *I*_Ks_ by shifting its activation curve towards positive potentials. This effect was not observed in cells transfected with the whole K_V_7.1/KCNE1 complex. These results suggest that, albeit obstructing RvD1 effect on the voltage dependence of K_V_7.1 gating, KCNE1 potentiates RvD1 inhibitory effect on the alpha channel subunit. It has been reported that K_V_ blocking agents show poor specificity because they mosty bind to conserved residues located in the pore domain (S6).^34^ However, KCNE1 regulates K_V_7.1 conductance by modulating the voltage sensor and its coupling with the pore domain.^35, 36^ Thus, we speculate that the β-subunit may modulate blockers interaction with the channel α-subunit allosterically; however, determining the RvD1 binding site on K_V_7.1/KCNE1 is beyond the scope of the present study. The observation that RvD1 increased the *I*_Ks_ step/tail ratio, indicating an effect inversely proportional to the current driving force, suggests that the block may be partially relieved by ions flowing through the channel.

Since RvD1 was more potent than LXA_4_ in blocking *I*_Ks_ in COS-7 cells, we focused on this SPM to analyze its effects on other cardiac K_V_ channels under heterologous expression. RvD1 effects on *I*_Kur_, *I*_to_ and *I*_K1_ was considerably smaller than on *I*_Ks_. A more substantial inhibitory effect was observed for *I*_Kr_ and found to be the most significant in prolonging repolarization in numerical simulations. The effect on *I*_Kr_ rectification (*I*_ss_/*I*_tail_ ratio) suggests RvD1 interference with the inactivation gating and might change the impact of *I*_Kr_ blockade on the AP profile (i.e. reduce the effect on phase 3 as compared to that on phase 2).

### 4.2. Pathophysiological relevance

RvD1 and LXA_4_ are enzymatically synthesized in response to inflammation during the resolution of the inflammation process; indeed, their levels increase during the resolution phase or inflammation.^37^ Serum levels of RvD1 of ≅ 160 pg/mL (0.34 nM) have been reported in humans.^38^ The present study tested ∼10 to 100-fold higher RvD1 concentrations; therefore the direct pathophysiological relevance of results may be questioned. Nonetheless, in local inflammation, tissue metabolites concentration might exceed by far circulating levels and RvD1 concentration is increased by blood coagulation,^39^ which is common in AF.

Inflammation has been described as a component of AF-induced atrial remodeling and contributes to atrial cardiomyopathy.^5,6^ Notably, the “metabolic syndrome” (encompassing hypertension, insulin resistance and obesity), a known risk factor for arrhythmogenesis, is characterized by a systemic inflammatory state.^2,7,11,40^ Recent studies have shown that RvD1 attenuates atrial fibrosis and reduces AF-inducibility, effects associated mainly to reversal of inflammation-induced structural and functional changes.^5,6^ Nonetheless, enhancement of potassium currents (e.g. *I*_Ks_) has been observed in cardiomyocytes from patients with chronic AF^41^ and RvD1 reversed atrial APD shortening in an ischemic heart failure model.^5^ Moreover, palmitic acid, another inflammatory lipid, increased *I*_Ks_ and *I*_Kr_ density in COS-7 cells and shortened APD in guinea-pig cardiomyocytes.^40^ Hence, direct inhibition of cardiac potassium channels, observed in the present work, might contribute to SPMs protective effect against arrhythmias, AF in particular.

A further possibility is that K^+^ currents inhibition by SPMs, possibly in immune cells, may contribute to their antiinflammatory action. Indeed, dofetilide a potent *I*_Kr_ blocker, has been reported to reduce the levels of IL-6, a cytokine involved in atrial inflammation.^42^ However, the relationship between inhibition of K^+^ currents and IL-production remains uncertain.

Using computational modeling, we examined the expected impact of RvD1 on atrial and ventricular repolarization. The results of these studies confirm that RvD1 effect on K^+^ currents may substantially prolong repolarization in both tissues. Albeit potentially beneficial in conditions in which shortened repolarization may contribute to arrhythmogenesis (e.g. AF), this effect might instead be of concern when ventricular repolarization is primarily prolonged, or repolarization reserve is reduced. The present results may thus be relevant to the design and use of SPMs-based agents, as part of the Comprehensive in Vitro Proarrhythmia Assay (CiPA) initiative.^43^

## 5. Conclusions

RvD1 directly inhibit a wide array of K^+^ currents relevant to myocardial repolarization. The findings are pathophysiologically relevant to conditions involving inflammatory and/or procoagulative states, the prerequisite for endogenous SPMs synthesis. It should be noted that K^+^ current inhibition may be beneficial under some conditions and undesirable in others.

## Supplementary Material

Supplementary material is available at Cardiovascular Research online.

## Authors’ contributions

C.V. and A.d.l.C. conceived the project, analyzed data, supervised the whole project. A.d.l.C., C.V. and A.Z. wrote the manuscript. A.d.l.C., P.G.S., A.d.B.-B. conducted and analyzed the experiments in cell lines under C.V. supervision. C.R. conducted and analyzed the experiments in guinea-pig cardiomyocytes under A.Z supervision. C.B. conducted the in silico experiments supervised by S.S. All authors have read and agreed to submit the present manuscript version.

## Acknowledgments

The authors are grateful to Dr Eva Delpón of the Universidad Complutense de Madrid, Spain for kindly providing us with K_ir_2.1 cloned into pcDNA3.1, to Drs S. Nattel, T. E. Hébert, and W. Weerapura of the Université de Montréal, Canada for kindly providing us with K_V_11.1 cloned into pcDNA3.1 and to Dr Michael M. Tamkun for providing us the stable cell line expressing K_V_1.5 channels. We also thank Dr. Lisardo Boscá for his helpful comments.

## Data availability

All raw data supporting the findings of this study are available from the corresponding authors upon reasonable request.

