## Supplemental for "RvD1 and LXA4 inhibitory effects on cardiac voltage-gated potassium channels"

[1] Instituto de Investigaciones Biomédicas Sols-Morreale CSIC-UAM, Madrid, Spain.
[2] Department of Biomedical and Clinical Sciences, Linköping University, Linköping, Sweden.
[3] Department of Biotechnology and Biosciences, Università degli Studi di Milano-Bicocca, Milan, Italy.
[4] Department of Electrical, Electronic and Information Engineering 'Guglielmo Marconi', University of Bologna, Cesena, Italy.
[5] Network of Cardiovascular Diseases (CIBERCV).

**Short title:** RvD1, $LXA_4$ and voltage-gated K channels

### This PDF includes the following figures and tables:

### Figures

Figure S1: Voltage-dependent effects of RvD1 on $K_V7.1$ in transiently transfected COS-7 cells

Figure S2: One-pulse RvD1 inhibition protocol on $I_{Ks}$ of guinea-pig cardiomyocytes

Figure S3: One-pulse $LXA_4$ inhibition protocol on $I_{Ks}$ of guinea-pig cardiomyocytes

Figure S4: RvD1 inhibition of $K_V11.1$ “tail current” traces and I-V relationship in the presence and the absence of RvD1 on $K_V11.1$

Figure S5: O’Hara–Rudy and Koivumaki computational modeling of RvD1 and $LXA_4$

Figure S6: RvD1 (5 and 50 nM) effects on the cardiac VG potassium channels

### Tables

Table S1. Computational modeling $APD_{90}$ results with the ORd model

Table S2. Computational modeling APD results with the atrial K model

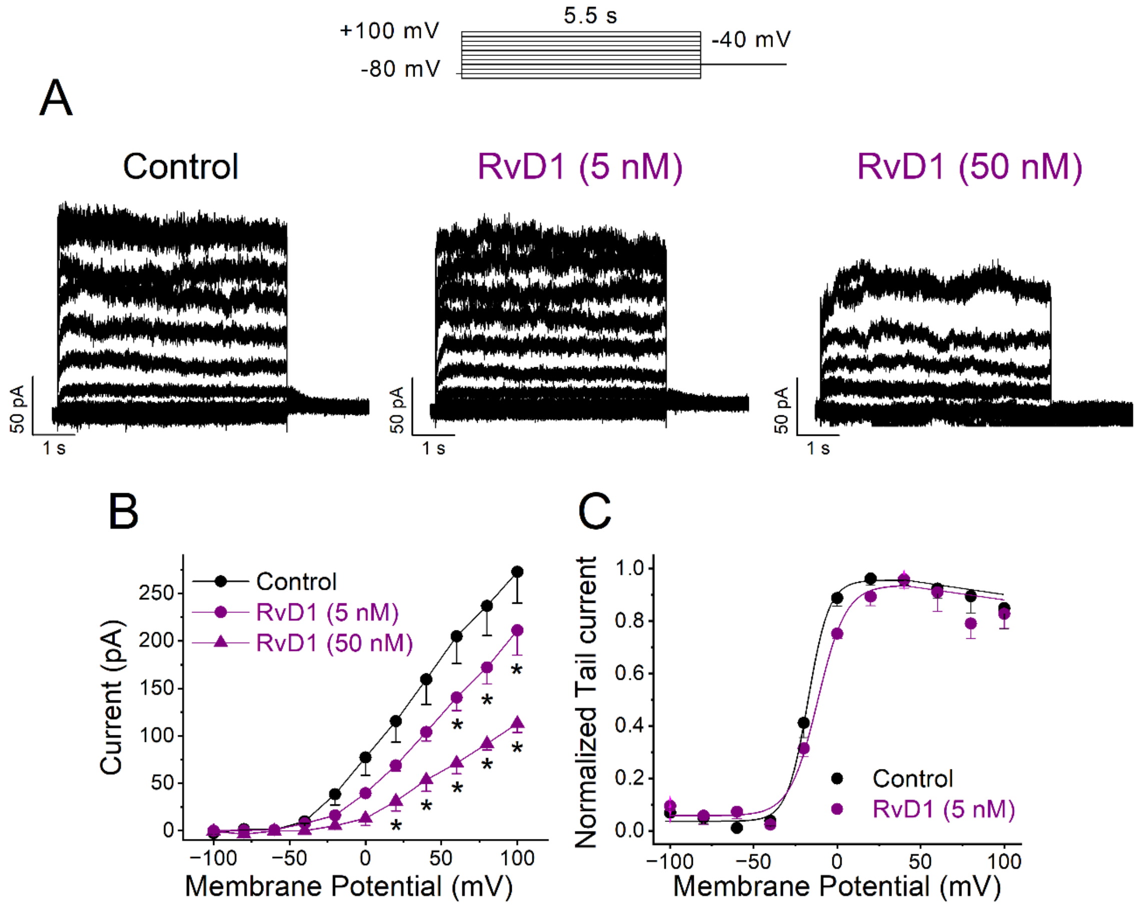


**Figure S1: Voltage-dependent effects of RvD1 on $K_V$7.1 in transiently transfected COS-7 cells**. A: Current traces obtained after applying the pulse protocol shown in the top, in the absence and in the presence of 5 and 50 nM RvD1. B: I-V relationships obtained after plotting the current at the end of 5.5-s vs. membrane potential, in the absence and in the presence of RvD1. C: Activation curves of $K_V$7.1 current obtained after representing the maximum tail current amplitude vs. the previous step potential, recorded in the absence and in the presence of 5 nM RvD1. Data are shown as the mean ± SEM. * $P < 0.05$ (Paired Student's t-test), n = 4-6.

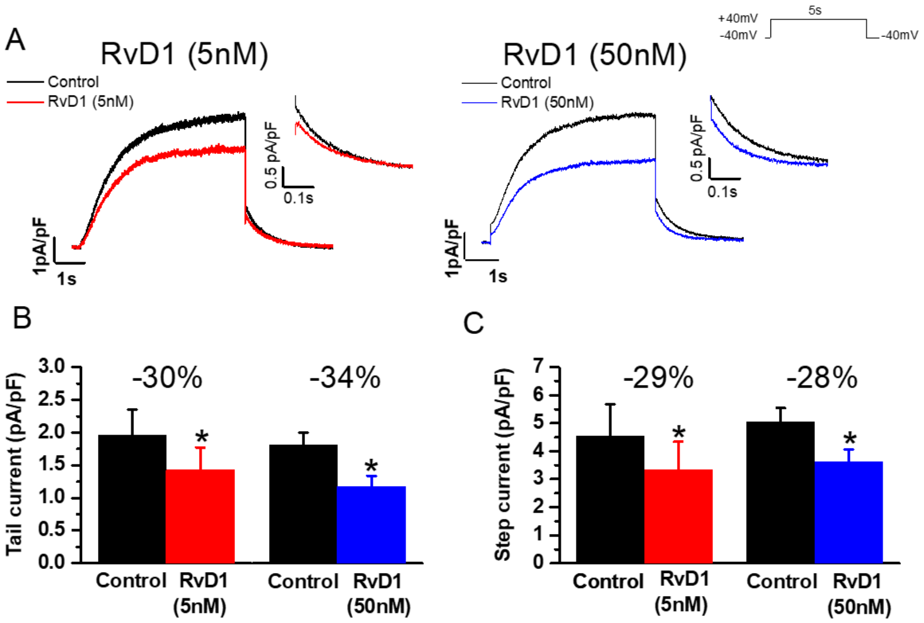


**Figure S2: One-pulse RvD1 inhibition protocol on $I_{Ks}$ of guinea-pig cardiomyocytes**. A: Representative traces of $I_{Ks}$ recorded within the same myocyte at +40 mV in control (black) and 5 nM RvD1 (red) or 50 nM RvD1 (blue); $I_{Ks}$ "tail current" magnified in the insets. B: Average (± SEM) density of "tail $I_{Ks}$" and C: $I_{Ks}$ at the end of the depolarizing step for all the treatments. n > 6 for all groups. * $P$ < 0.05 vs. control (Paired Student's *t-test*).

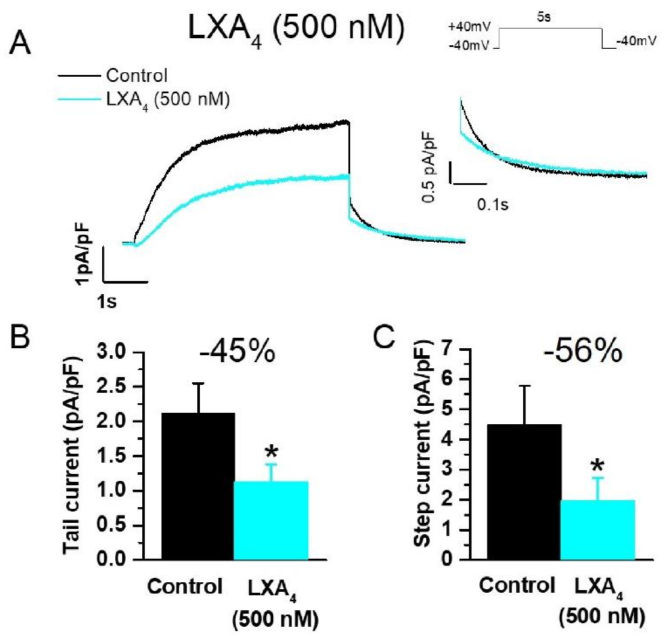


**Figure S3: One-pulse $LXA_4$ inhibition protocol on $I_{Ks}$ of guinea-pig cardiomyocytes**. A: Representative traces of $I_{Ks}$ recorded within the same myocyte at +40 mV in control (black) and in the presence of 500 nM $LXA_4$ (light blue); $I_{Ks}$ "tail current" magnified in the inset. B: Average (± SEM) density of "tail $I_{Ks}$" and C: $I_{Ks}$ at the end of the depolarizing step. n = 8 for all groups. * $P < 0.05$ vs. Control (Paired Student's t-test).

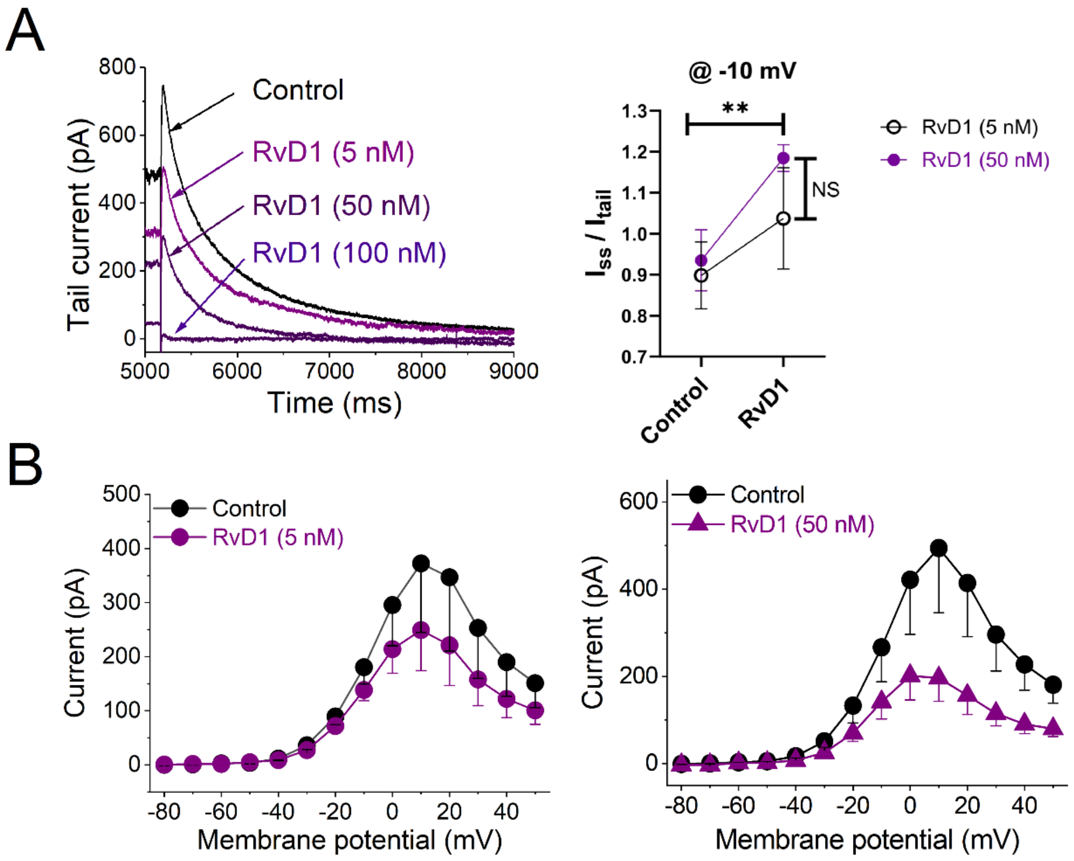


**Figure S4: RvD1 inhibition of $K_V$11.1 "tail current" traces and I-V relationship in the presence and the absence of RvD1 on $K_V$11.1**. A: Left panel: $K_V$11.1 "tail current" traces in the absence and the presence of 5 nM, 50 nM and 100 nM of RvD1 measured at +20 mV membrane potential. Right panel: Effect of RvD1 at 5 and 50 nM on $I_{ss}/I_{tail}$ measured at -10 mV. B: I-V relationship of $K_V$11.1 current in the absence and the presence of 5 nM RvD1 (left panel) and 50 nM RvD1 (right panel) measured at the end of the 5 s pulse and plotted vs. membrane potential. Data are shown as the mean ± SEM. * $P < 0.05$ (Paired Student's t-test), n = 4-6.

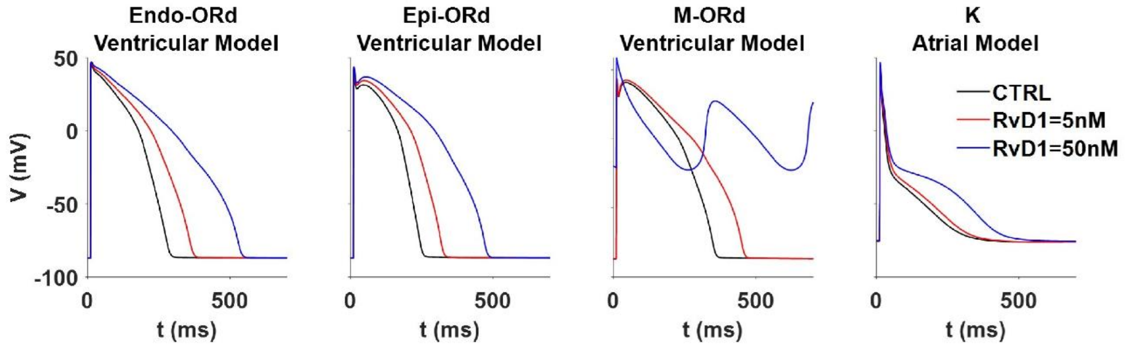


**Figure S5: O'Hara–Rudy and Koivumaki computational modeling of RvD1**. ORd ventricular cell computer model was used to examine the effects of RvD1, on AP behaviors in A: endocardial, B: epicardial, and C: M cells. D: Model dependency test by simulating the effects with the K model of the effects of RvD1, on atrial AP morphology.

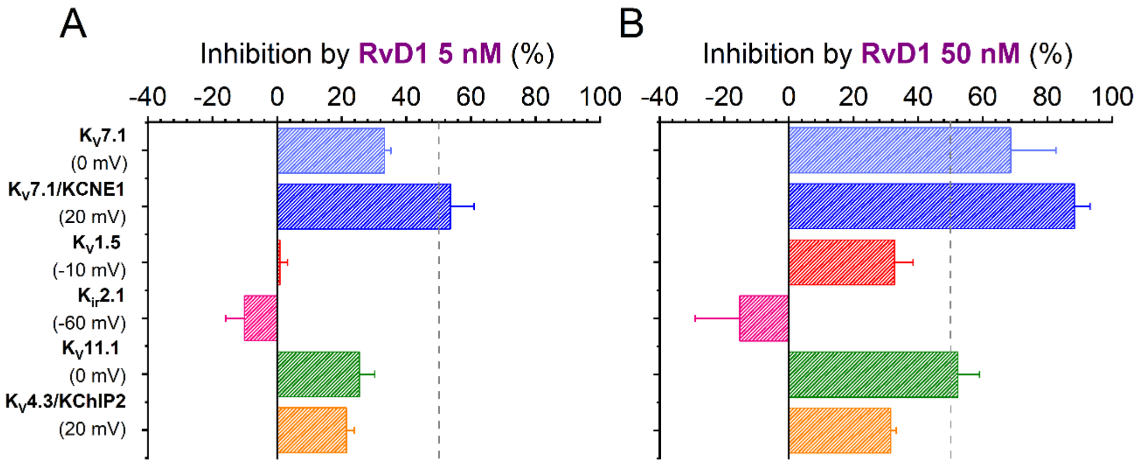


**Figure S6: RvD1 (5 and 50 nM) effects on the cardiac VG potassium channels**. A: RvD1(5 nM) effects on the different ion channels plotted vs. the percentage of inhibition measured at each $V_{1/2}$ for each ion channel. B: RvD1(50 nM) effects on the different ion channels plotted vs. the percentage of inhibition measured at each $V_{1/2}$ for each ion channel.

Table S1. Computational modeling $APD_{90}$ results with the ORd model

| | *$APD_{90}$ (ms)* | | | | | |
|---|---|---|---|---|---|---|
| | *Endo* | *Epi* | *M* | *Δ% Endo* | *Δ% Epi* | *Δ% M* |
| *CTRL* | 265 | 231 | 336 | | | |
| *RvD1 (5 nM)* | 346 | 305 | 436 | 31 | 32 | 30 |
| *RvD1 (5 nM) only $I_{Ks}$* | 272 | 237 | 350 | 3 | 3 | 4 |
| *RvD1 (5 nM) only $I_{to}$* | 264 | 231 | 337 | -0.4 | 0 | 0.3 |
| *RvD1 (5 nM) only $I_{Kr}$* | 330 | 289 | 406 | 25 | 25 | 21 |
| *RvD1 (50 nM)* | 513 | 455 | Rep failure | 94 | 97 | |
| *RvD1 (50 nM) only $I_{Ks}$* | 279 | 243 | 363 | 5 | 5 | 8 |
| *RvD1 (50 nM) only $I_{to}$* | 264 | 231 | 338 | -0.4 | 0 | 1 |
| *RvD1 (50 nM) only $I_{Kr}$* | 442 | 387 | 555 | 67 | 68 | 65 |

Table S2. Computational modeling APD results with the atrial K model

| | *APD (ms)* | *Δ%* |
|---|---|---|
| *CTRL* | 233 | |
| *RvD1 (5 nM)* | 259 | 11 |
| *RvD1 (5 nM) only $I_{Ks}$* | 238 | 2 |
| *RvD1 (5 nM) only $I_{to}$* | 257 | 10 |
| *RvD1 (5 nM) only $I_{Kr}$* | 239 | 3 |
| *RvD1 (5 nM) only $I_{Kur}$* | 225 | -3 |
| *RvD1 (50 nM)* | 370 | 59 |
| *RvD1 (50 nM) only $I_{Ks}$* | 241 | 3 |
| *RvD1 (50 nM) only $I_{to}$* | 257 | 10 |
| *RvD1 (50 nM) only $I_{Kr}$* | 247 | 6 |
| *RvD1 (50 nM) only $I_{Kur}$* | 263 | 13 |
